# Subchondral bone marrow adipose tissue lipolysis regulates bone formation in hand osteoarthritis

**DOI:** 10.1101/2024.09.26.615232

**Authors:** Mauro Maniglio, Léa Loisay, Diego de Haro, Alexander Antoniadis, Thomas Hügle, Jeroen Geurts

## Abstract

**Objective:** Bone marrow adipose tissue (BMAT) is emerging as an important regulator of bone formation and energy metabolism. Lipolysis of BMAT releases glycerol and fatty acid substrates that are catabolized by osteoblasts. Here, we investigated whether BMAT lipolysis is involved in subchondral bone formation in hand osteoarthritis (OA).

**Methods:** Subchondral BMAT lipolysis and bone marrow adipocyte (BMAd) morphology were studied in clinical specimens of carpo-metacarpal (CMC1) ans distal interphalangeal joint (DIP) OA. BMAd size, osteoblast numbers and expression of lipolysis enzymes (ATGL, phospho-HSL, MGLL) were compared between regions of low and high bone formation. Free fatty acids, glycerol and bone biomarkers were measured in osteochondral explants.

**Results:** Subchondral BMAd size was positively correlated with BMI and reduced in regions of high bone formation. Osteoblast numbers were negatively correlated with BMAd size. ATGL, phoshpo-HSL and MGLL were expressed in both in BMAds and activated osteoblasts and increased in regions of high bone formation. Secreted glycerol levels, but not free fatty acids, were correlated with bone formation markers pro-collagen type I and alkaline phosphatase.

**Conclusion:** Our findings reveal a previously unrecognized role of BMAT lipolysis in regulating bone formation in hand OA, which may be modulated by BMI.

## Introduction

Bone marrow adipose tissue (BMAT) is one of the major fat depots and accounts for approximately ten percent of total fat mass in lean, healthy humans. The notion that BMAT serves merely as an inert filler of the bone cavity has evolved over the last decade and this tissue is now considered an important regulator of bone homeostasis and systemic energy metabolism [1-3]. BMAT is dynamic tissue and its composition and volume undergo changes in response to states of caloric excess and deprivation [4, 5]. BMAT volume increases with ageing, obesity, caloric restriction, diabetes, and osteoporosis, while decreasing by cold exposure, exercise and gastrectomy [6, 7]. As bone marrow adipocytes (BMAds), osteoblasts and hematopoietic cells share their location within the bone cavity, reduction in BMAT volume is associated with bone accrual [8].

BMAds are specialized for the storage of energy in the form of neutral lipids such as triglycerides and sterol esters. Triglycerides are broken down and released in states of energy demand by a catabolic process termed lipolysis involving the sequential activities of three enzymes: adipose triglyceride lipase (ATGL encoded by the gene *PNPLA2*), hormone-sensitive lipase (HSL encoded by the gene *LIPE*) and monoglyceride lipase (MGLL encoded by the gene *MGLL*) [9]. Lipolysis yields free glycerol and free fatty acids (FFA) as energy substrates, which have been described to fuel bone formation. Mitochondrial beta-oxidation of free fatty acids in osteoblasts is an essential mechanism for postnatal bone acquisition [10]. Adipocyte-specific knockout of *Pnpla2* reduced parathyroid hormone-induced bone formation [11] and BMAd-specific *Pnpla2* knockout animals displayed reduced *de novo* bone formation in cortical bone defects and bone loss in response to chronic cold exposure [6]. Osteoblast-specific *Pnpla2* deficiency reduced both cortical and trabecular bone parameters and led to an accumulation of triglycerides in bone [12]. Notably, ablation of adiponectin-expressing cells in BMAT yielded the most rapid and profound increase in systemic bone mass yet observed [8]. Therefore, the ability of BMAT to modulate its surrounding bone tissue indicates that this adipose depot could be a novel regulator of subchondral bone remodeling in OA.

Hand OA frequently affects the thumb base (CMC-1) and interphalangeal (IP) joints and is characterized by subchondral bone sclerosis, bone marrow lesions, cysts and osteophytes [13]. Hand OA is highly associated with high BMI and obesity [14-16] and several adipose-derived systemic mediators including adiponectin, leptin, cholesterol and fatty acids have been linked to an increased risk for hand OA [13, 17, 18]. However, it remains unknown whether subchondral BMAT in hand OA is affected by obesity and contributes to pathological features in bone.

Here, we investigated whether BMAT lipolysis is involved in subchondral bone formation in hand OA. Clinical hand OA specimens showed a specific reduction of BMAd size and increased expression of ATGL, phosphorylated HSL and MGLL in regions of active bone formation. BMAd size was correlated with BMI. Employing an explant model, we found a correlation between secreted glycerol levels and bone biomarkers. Our findings reveal an important role of BMAT lipolysis in regulating bone formation in hand OA, which may be modulated by BMI.

## Methods

### Patients and sample collection

The study protocol was approved by the local ethic committee (CER-VD 2024-0117). Clinical specimens were obtained through convenience sampling and experiments were performed with ≥10 donors, based on our previous work [19, 20], to obtain biologically significant results. Waste material was acquired from patients (*n* = 23, 14 female, mean age 63 years, range 43–92 years) undergoing surgery for advanced symptomatic CMC-1 OA (*n* = 19) or distal IP OA (*n* = 4). Specimens were collected immediately in 10% formalin buffer for histological processing or PBS for explant culture. Fresh samples were processed within 2 hours after surgery. Patient characteristics are shown in Table 1. Tibia plateaus (*n* = 5) were obtained from knee arthroplasty surgery and used for isolation of primary osteoblasts.

**Table 1.**
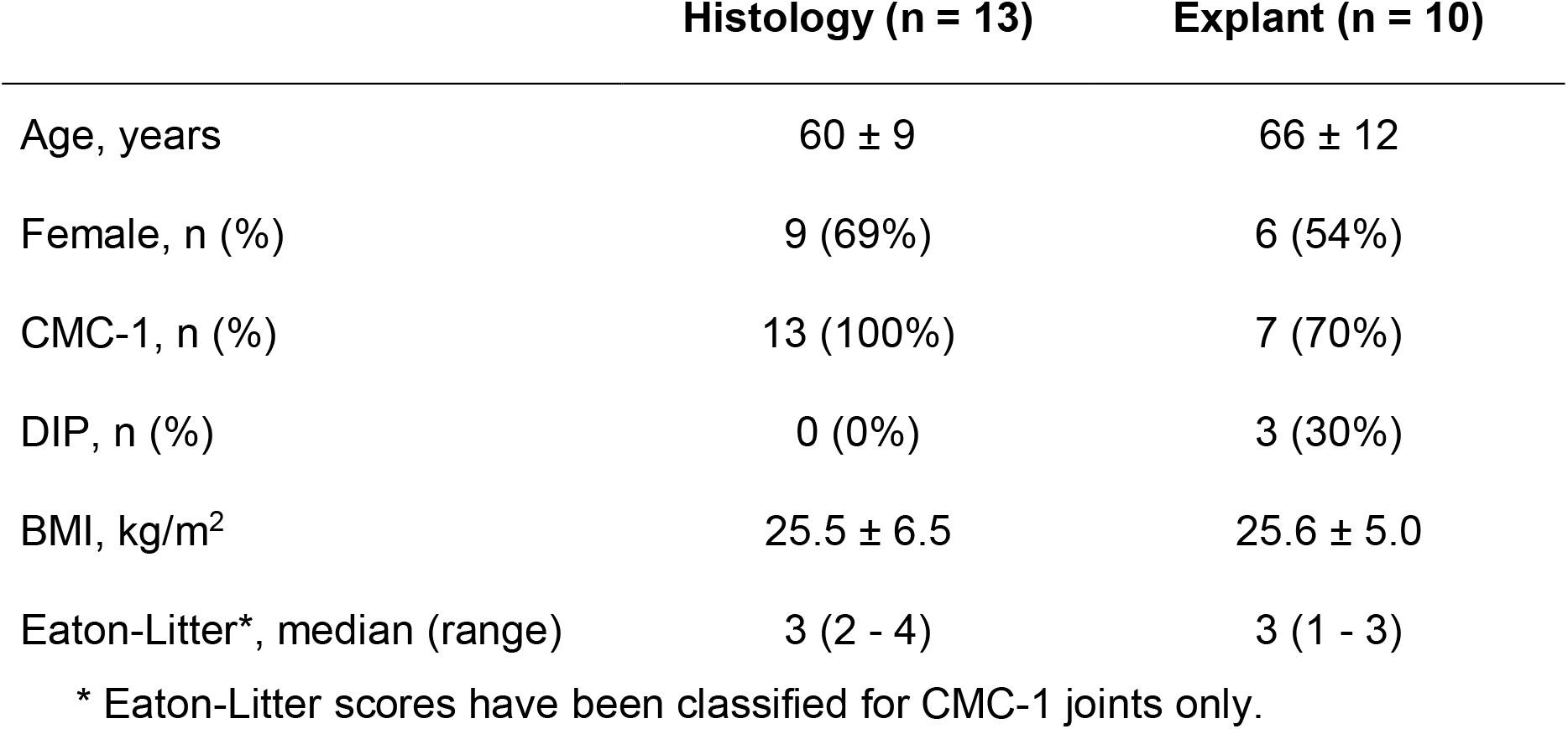
Descriptive characteristics of clinical specimens.

### Radiological analysis

Radiological CMC-1 OA was graded using the Eaton-Littler classification on lateral radiographs [21]. Stage 1: Subtle CMC joint widening. Stage 2: Slight CMC joint space narrowing, sclerosis, cystic changes and osteophytes. Stage 3: Advanced CMC joint space narrowing, sclerosis, cystic changes and osteophytes. Stage 4: Scaphotrapezial OA and advanced CMC joint space narrowing, sclerosis, cystic changes and osteophytes.

### Whole-mount staining

Fixed specimens were rinsed in PBS and stained with 0.3% Oil red O solution (#01391, Sigma-Aldrich) in 60% isopropanol, washed briefly with PBS and counterstained with Hoechst staining (#33342, 1 μg/mL, Sigma-Aldrich). Images were acquired on a Leica M205-FA stereomicroscope with LAS X software.

### Histological analysis

Specimens were fixed in 10% neutral-buffered formalin for 2 days, decalcified in 5% formic acid and paraffin-sectioned at 5 μm thickness. Sections were stained with Safranin-*O* and fast green and RGB trichrome according to standard protocols [22]. Immunohistochemistry was performed on serial sections as previously described [23] using the following primary antibodies: anti–MGLL (1:100; #HPA011993, Sigma-Aldrich), anti–HSL (1:100; #4107, Cell Signaling Technology), anti–phospho-HSL (1:100; #4139, Cell Signaling Technology), anti-ATGL (1:100; #2138, Cell Signaling Technology), anti-osteocalcin (1:200; MAB1419, R&D Systems). Immunoreactivities were visualized using mouse or rabbit-sepcific Vectastain Elite ABC kit (Vector Laboratories). Negative controls were performed with both isotype controls and secondary antibodies only (Figure S1). Sections were counterstained with papanicolaou staining. For anti-HSL, anti-phospo-HSL and anti-ATGL immunostaining, antigen retrieval was performed in citrate buffer (pH 6.0) at 60 °C overnight. Images were acquired on a NanoZoomer S60 Digital slide scanner (Hamamatsu Photonics).

### BMAT image analysis

BMAd size was quantified from slide-scanned images of Safranin-*O*-stained tissue sections using ImageJ. ROIs displaying high bone remodeling were selected based on bone area fraction >50%, BMAT/total marrow fraction <80% and/or presence of BMAT fibrosis on RGB-stained sections (Figure S2A). ROIs displaying low bone remodeling were selected based on bone area fraction <50%, BMAT/total marrow fraction >80% and absence of BMAT fibrosis. Greyscale images were thresholded using the Huang method, followed by segmentation using a watershed algorithm [24]. A minimum of 200 and 25 BMAds were quantified in respectively low-power and high-power fields (HPF) subjacent to the subchondral bone plate. The number of osteoblasts were counted manually in HPF. Quantification of IHC staining was performed by image binarization, followed by pixel counting using Adobe Photoshop. All quantifications were performed unblinded.

### Explant culture

CMC-1 or IP bone explants were cultured in Krebs-Ringer buffer (Sigma-Aldrich# K4002) supplemented with 150 mM NaHCO_3_, 10 mM HEPES and 0.5% BSA at 37°, 5% CO_2_ during 2 days [25]. Supernatant was collected and stored at -80°C until further analysis. Explants were fixed in 10% neutral-buffered formalin.

### Primary osteoblast culture

Primary human osteoblasts were obtained by outgrowth culture from bone chips from tibia plateaus as described previously [19]. First passage osteoblasts were cultured in osteogenic medium (DMEM, 10% FBS, 200 mM L-glutamine, 50 µg/mL ascorbic acid, 2mM β-glycerophosphate) for 7 days.

### Gene expression analysis

RNA was extracted using TRIZOL reagent according to standard protocols. RT-qPCR was performed using PowerUp SYBR Green (#A25742, Applied Biosystems) using a QuantStudio 5 Real Time PCR system (Applied Biosystems). Primers are listed in Table S1. Relative gene expression was quantified using ΔCt method and *GAPDH* as reference gene.

### Supernatant analysis

Adiponectin (#DY1065-05, R&D Systems) and pro-collagen type I (#DY6220-05, R&D Systems) were measured using ELISA kits. Glycerol (#MAK266, Sigma-Aldrich) and free fatty acids (#MAK044, Sigma-Aldrich) were assessed using colorimetric assays. Alkaline phosphate activity was assayed using p-nitrophenyl phosphate ((#S0942, Sigma-Aldrich) in alkaline buffer solution (pH 10.3, #A9226, Sigma-Aldrich) for 2 hours at 37 °C. All quantifications were normalized to explant weight.

### Statistical methods

All data points on graphs indicate individual donors. Statistical tests were performed using GraphPad Prism v9. The tests used to determine statistical significance are indicated in figure legends. Variables from all experiments were assessed for normality with Anderson-Daarling, Shapiro-Wilk, Kolmogorov and D’Agostino-Pearson tests. Data were analysed using paired *t*-test or Wilcoxon matched-pairs signed-rank test. Correlation analysis was performed using Pearson or Spearman correlation.

## Results

### BMAd hypertrophy is dependent of BMI in CMC-1 OA

To investigate whether BMI affects BMAT properties in CMC-1 joints, we analysed BMAd size in ROIs of low bone remodeling. Average BMI was 25.5 ± 6.5 and ranged from 19– 42 kg/m^2^. Histological analyses revealed similar BMAT area fraction (Figure 1A), while BMAd size distribution was changed between specimens (Figure 1B). The proportion of BMAds smaller than 400 μm^2^ declined (30%, 17%, 10% for BMI 19, 27, 43 respectively), whereas the proportion of BMAds larger than 1000 μm^2^ increased with increasing BMI (9%, 43%, 55% for BMI 19, 27, 43 respectively). Average BMAd size was positively correlated with BMI (Pearson *r* = 0.60, p = 0.02; Figure 1C). These findings reveal that BMAd hypertrophy in the CMC-1 joint may be dependent on BMI and obesity.

**Figure 1.**
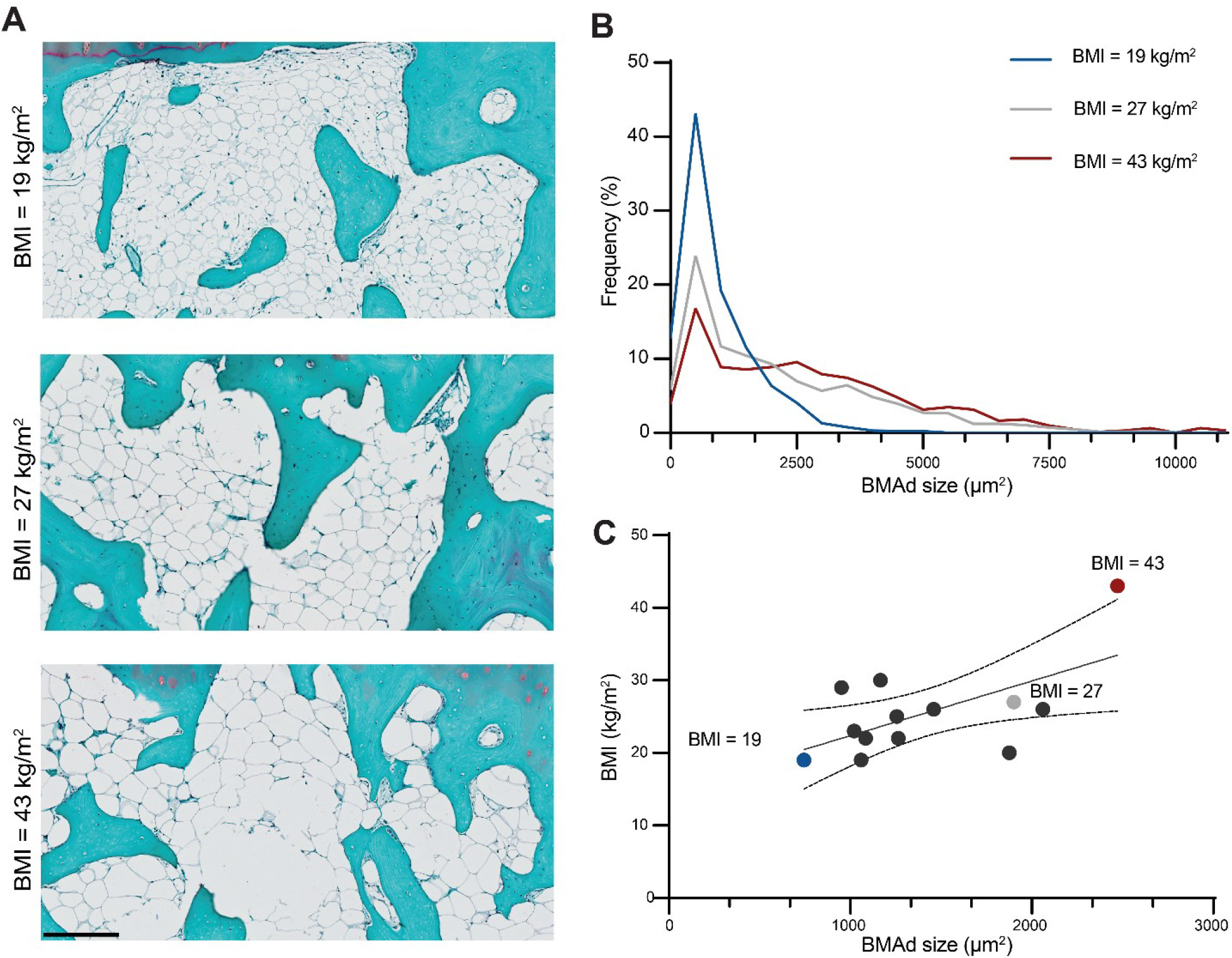
BMAd size distribution in CMC-1 joint of patients with different BMI. Typical morphology of subchondral BMAT visualized by Safranin-O staining. Scale bar = 200 μm (A) Frequency distribution of BMAd size in three samples from patients with the lowest (blue, 19 kg/m^2^), average (grey, 27 kg/m^2^) and highest BMI (red, 42 kg/m^2^). (B) Scatterplot showing correlation between BMI and median BMAd size in 13 CMC-1 joints and the regression line with 95% confidence intervals. (C)

### BMAd size decreases in regions of bone remodeling

To understand whether bone formation in CMC-1 OA is regulated by BMAT changes, we compared BMAd size and its correlation with osteoblast numbers in regions of low and high bone formation within specimens. Whole-mount lipid staining revealed distinct BMAT morphologies between these regions, characterized by less intense lipid staining, smaller BMAds and increased cell numbers detected by Hoechst staining in high remodeling regions (Figure 2A). These findings were confirmed by histological analysis, revealing smaller BMAds, increased cell numbers and extensive fibrosis in BMAT of high remodeling regions (Figure 2B). Analysis of average BMAd size revealed an approximately two-fold reduction between regions of low and high remodeling (p<0.001); Figure 2C). Osteoblast numbers were significantly increased in high remodeling regions (p<0.001; Figure 2D). Average BMAd size was negatively correlated with osteoblast numbers (Pearson *r* = –0.48, p = 0.01; Figure 2E). Together, these findings reveal that BMAT undergoes extensive morphological changes during bone remodeling and that BMAd size is associated with activity of bone-forming cells.

**Figure 2.**
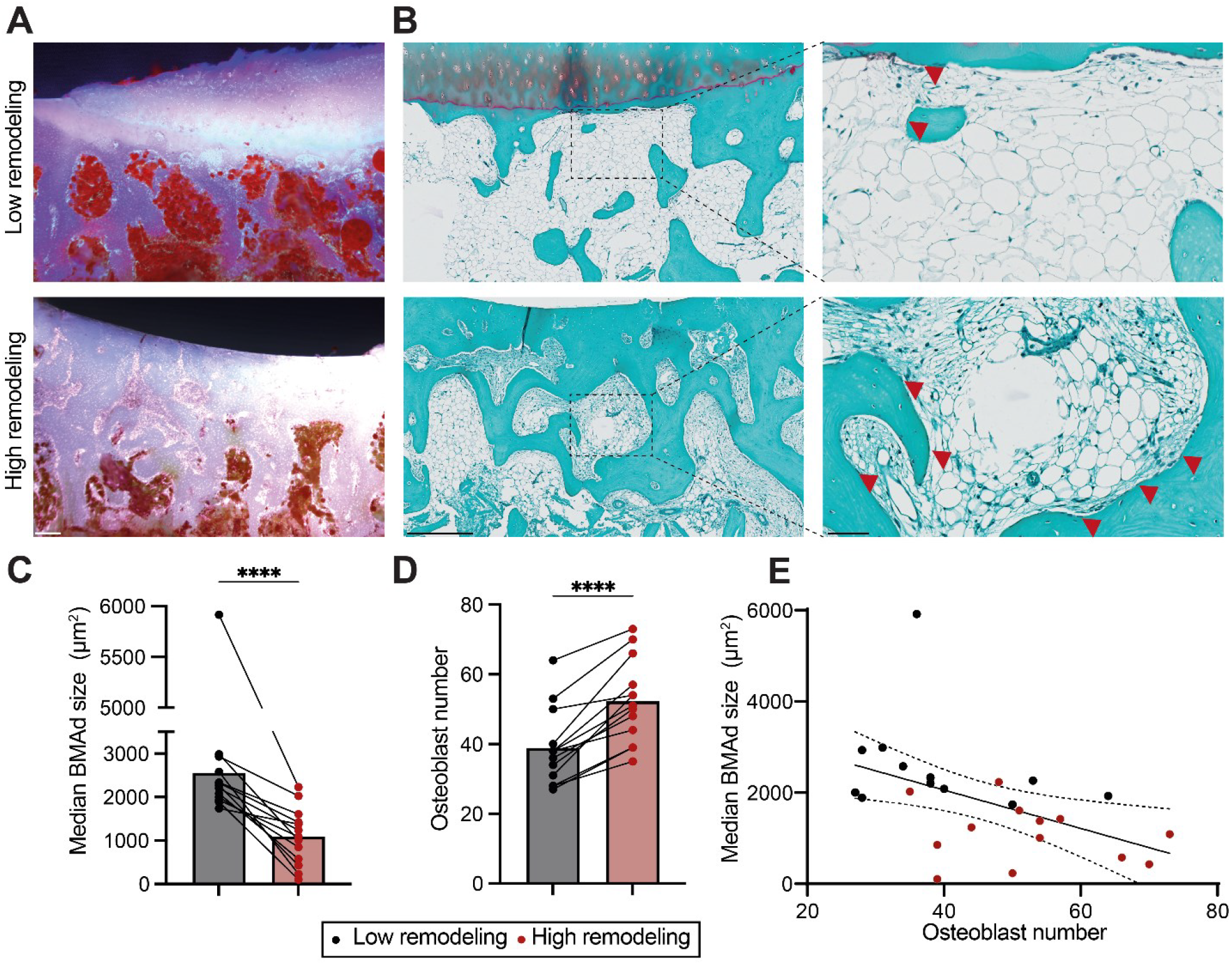
Reduction of BMAd size in areas of high bone remodeling. Typical morphology of subchondral bone and BMAT in low and high remodeling areas visualized by whole-mount Oil red-O and Hoechst staining. Scale bar = 200 μm. (A) Typical morphology of subchondral BMAT in areas of low and high remodeling visualized by Safranin-O staining. Osteoblast indicated by red arrowheads. Scale bars = 500 μm (low magnification) and 100 μm (high magnification). (B) Median BMAd size quantified from high magnification images. P-value < 0.001 by paired *t*-test. (C) Osteoblast numbers quantified from high magnification images. P-value < 0.001 by paired *t*-test. (D). Scatterplot showing correlation between median BMAd size and osteoblast numbers in low and high remodeling areas of 13 CMC-1 joint and the regression line with 95% confidence intervals. (E)

### Increased BMAT lipolysis in CMC-1 OA

Adipocyte size is regulated by the rates of lipogenesis and lipolysis [8]. To determine whether reduced BMAd size may be caused by increased lipolysis, we performed IHC analyses of the three enzymes involved in this process. Expression of ATGL, the rate-limiting enzyme of lipolysis, was increased 1.67-fold (p = 0.006) in BMAT of regions with high bone remodeling (Figure 3A). Interestingly, ATGL was expressed in both BMAds and activated bone-lining cells. While HSL was expressed primarily in BMAds without differences between ROIs (Supplementary Figure), the activated form phospho-HSL was increased 1.90-fold (p = 0.004) in regions of high remodeling (Figure 3B). MGLL expression was increased 3.48-fold (p = 0.0005) and localized to both BMAds and activated bone-lining cells. Lipolysis genes *PNPLA2, LIPE and MGLL* were expressed by primary human osteoblasts (Figure 3E). Overall, these data suggest that BMAT lipolysis contributes to subchondral bone formation in CMC-1 OA.

**Figure 3.**
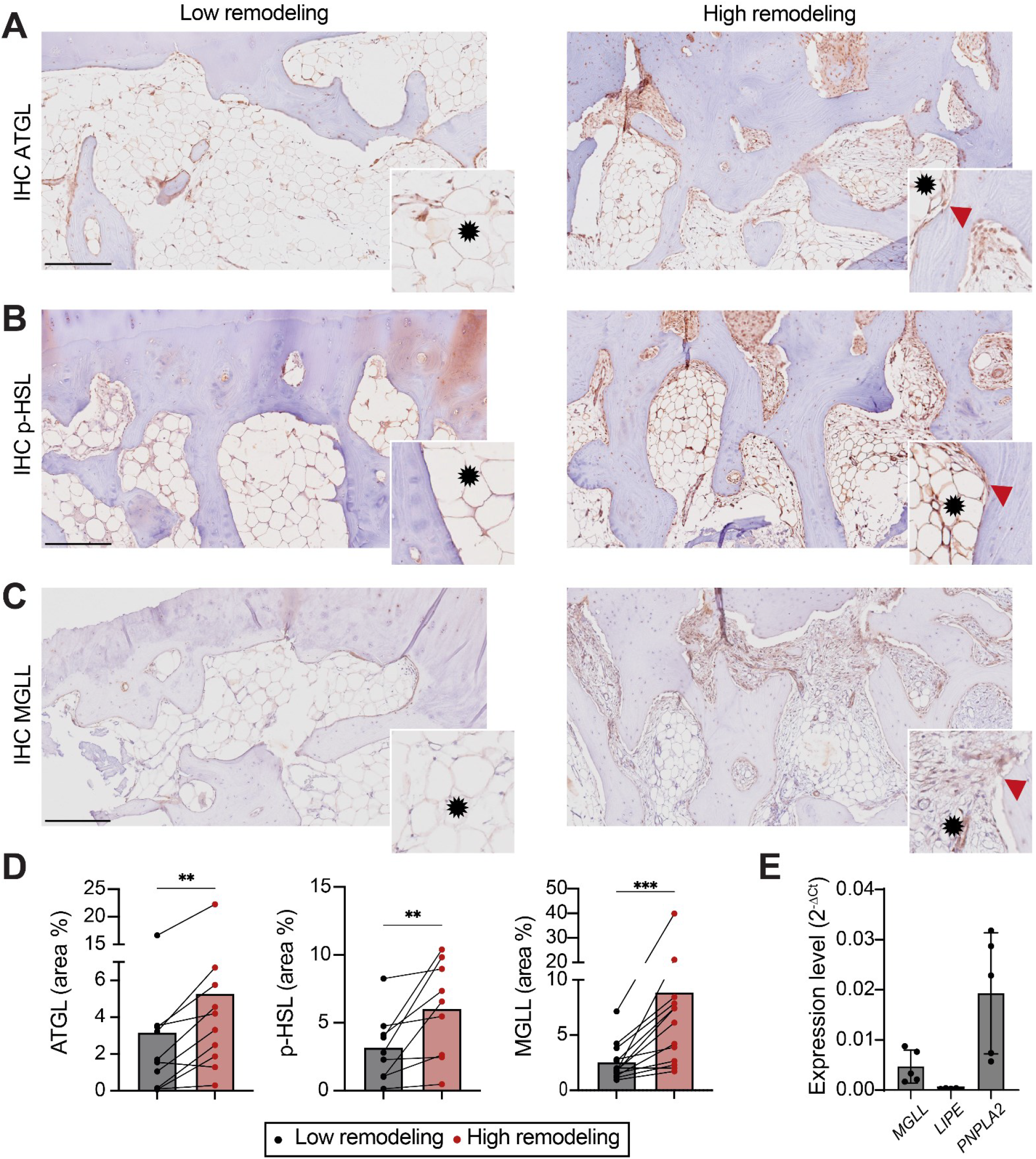
Increased expression of lipolysis enzymes in high remodeling areas. Representative images of ATGL (A), phospho-HSL (B) and MGLL (C) immunostaining in low and high remodeling subchondral BMAT in CMC-1 joints. Scale bar = 250 μm. Image insets indicate positive immunostaining in BMAd (asterisk) and bone-lining cells (red arrowhead). (A-C) ATGL-(*n* = 10), phospho-HSL-(*n* = 9) and MGLL-positive (*n* = 13) area fraction quantified from IHC images. P-value < 0.01 by Wilcoxon matched-pair signed rank test. (D) Expression levels of lipolysis genes *MGLL, LIPE, PNPLA* in OA-derived osteoblast from five knee joints after 7 days of culture in osteogenic medium.

### Lipolysis end product glycerol correlates with bone formation markers

To study whether lipolysis plays a role in fueling bone formation in hand OA, we measured bone formation markers and lipolysis end products, glycerol and FFA, in bone explants of CMC-1 and distal IP OA joints. Median secretion levels were FFA: 0.85 (0.42-2.4) pmol/mg, glycerol: 10.2 (0.24–27.4) pmol/mg, adiponectin: 94.9 (16.9–214.9) ng/mg, pro-collagen type I: 34.4 (21.2–114.1) ng/mg and ALP 8.42 (7.55–8.55) mU/mg. Glycerol, but not FFA, was significantly correlated with pro-collagen type I, ALP (Table 2).

**Table 2.** Spearman correlation coefficients between bone markers and BMAT-derived mediators in explanted CMC-1 and DIP joints (*n* = 10).

|  | ALP | Pro-collagen type I |
| --- | --- | --- |
| Glycerol, *r* (*p*-value) | 0.76 (**0.037**) | 0.90 (**0.001**) |
| FFA, *r* (*p*-value) | 0.10 (0.840) | 0.38 (0.279) |
| Adiponectin, *r* (*p*-value) | 0.55 (0.171) | 0.61 (0.067) |
| BMI, *r* (*p*-value) | – 0.20 (0.641) | 0.05 (0.896) |
| ALP, *r* (*p*-value) |  | 0.60 (0.132) |

## Discussion

BMAT has emerged as a key regulator of bone homeostasis and energy metabolism. However, whether its reduction in subchondral bone plays a role in the context of OA remains unclear. The rapid and extensive bone formation observed following the ablation of BMAds in mice [8] shows striking histological similarities with subchondral bone sclerosis observed in advanced knee OA [19, 26]. This prompted us to focus on this understudied joint tissue in our research. In this study, we report that subchondral BMAd size is reduced in regions of high bone remodeling and formation in CMC-1 OA. This observation led us to investigate BMAT lipolysis through histological and biochemical analyses. These analyses revealed increased expression of lipolysis enzymes in both BMAds and active bone-lining cells, as well as a correlation between glycerol, the end product of lipolysis, and bone formation markers. Notably, BMAd size in CMC-1 joints appeared influenced by BMI. Together, our findings suggests that BMAT play a previously unrecognized role in regulating hand OA and may offer new insights into the link between OA risk due, obesity and dyslipidemia.

The association of obesity and an increased risk of OA in both non-weight-bearing and weight-bearing joints is thought to result from a combination of mechanical and metabolic effects [27]. Systemic mediators derived from adipose tissues, including fatty acids and adipokines, have been linked to structural and clinical features of hand OA. Low serum adiponectin levels [14] and postprandial saturated and polyunsaturated fatty acids levels [17] associated with higher hand OA risk after adjusting for total fat mass or BMI. Whether these systemic mediators can directly alter bone turnover and remodeling remains unknown. Modulation of intra- and periarticular fat depots in obese individuals may represent an additional mechanism through which obesity contributes to OA risk. The properties of the infrapatellar fat pad in particular, have been extensively investigated in the context of knee OA [28-31]. While histological studies have not shown BMI-dependent adipocyte hypertropy or necrosis in knee OA [28], fat pad hyperplasia appeared a better predictor of OA progression than BMI changes [30]. Our findings suggest that BMAd hypertropy may be BMI-dependent in the CMC-1 joint, an effect which could be explained by BMAT accumulation in response to caloric overfeeding in obese individuals. Interestingly, BMAT changes induced by high calorie feeding showed opposing effects at vertebral and femoral regions and higher BMAT accumulation in women [5]. Additionally, BMAds undergo distinct transcriptomic and lipidomic changes in states of caloric deficit or excess, both of which likely affect OA-related bone remodeling processes.

A major finding is that BMAT lipolysis is elevated in subchondral bone of CMC-1 joints and glycerol levels were correlated with markers of bone formation in explants of CMC-1 and DIP joints. Given that glycerol and free fatty acids are used in respectively gluconeogenesis and beta-oxidation, our findings remain inconclusive as to which pathway could be primarily fueling OA-related bone formation. Indeed, aerobic glycolysis and oxidative phosphorylation are both known to play a role in osteoblast maturation and regulation of bone mass [10, 32, 33] and differential utilization of these pathways is likely sex-specific and regulated by nutrient availability and metabolic needs of activated bone cells. Supporting this, deletion of ATGL in either BMAT [6] or all fat tissues [11] did not result in any changes in bone volume fraction. Osteoblast-specific deletion of ATGL led to a modest (∼10%) reduction of bone volume fraction in male, but not female mice. Notably, mature osteoblasts rely more on oxidative phosphorylation than undifferentiated stromal cells [34]. While lipolysis appears to play a limited role in general bone homeostasis, ATGL deficient mice exhibit significantly reduced bone formation rates during anabolic therapy [11] and delayed healing of cortical bone defects [6]. Given the importance of mechanical loading in the progression of OA, it is notable that BMAT lipolysis has been described to be induced through mechanosensitive factors, such as reticulocalbin-2 (RCN2) [35]. We hypothesize that both mechanical and metabolic factors could converge on subchondral BMAT lipolysis as a shared mechanism driving bone changes in hand OA.

The therapeutic implications of our findings are yet to be determined, as we currently lack specific inhibitors of lipolysis enzymes in humans. Systemic lipid species have been primarily under investigation as biomarkers [17, 36], although a number of retrospective cohort studies have evaluated therapeutic effects of targeting lipids directly or indirectly in hand OA. Statin therapy, which lowers cholesterol and triglyceride levels, proved ineffective in reducing the risk of incident hand OA in a sequential cohort study [37]. Hormone replacement therapy, when started around the onset of menopause, led to a decreased risk of hand OA, yet protective effects disappeared upon treatment cessation [38]. Interestingly, treatment with testosterone, but not estrogen, lowers glycerol levels and HSL enzyme activity in post-menopausal women [39]. Additionally, lower sex hormone levels have been linked to higher BMAT fraction in older men and women [40]. Together, there appears to be merit in further evaluating the role of BMAT lipolysis in the context of both pathology and treatment of hand OA.

We acknowledge several limitations of our study. First, this is a cross-sectional, within-patient comparative histological analysis of a relatively small joint, which presents challenges in distinguishing between remodeling and non-remodeling areas. Additionally, clinical specimens were not collected using a standardized method, which may introduce variability between samples. We sought to address these issues by establishing criteria for selecting ROIs and applying within-subject statistics. Second, we do not have additional data (e.g. histology) available to stratify samples used for explant studies, nor could we perform mechanistic studies on lipolysis in this setting. We are currently performing studies using explants from larger joints to establish correlations between histological and molecular readouts of BMAT lipolysis.

In summary, this study is the first to dissect subchondral BMAT lipolysis and identifies BMAT as previously unrecognized regulator of bone formation in in the context of hand OA.

## Acknowledgements

The authors thank Prof. Wassim Raffoul for his support. We are grateful to Elke Moradpour for her expert support in microscopy and histology.

## Author contributions

MM: Conceptualization, experimental design, data acquisition, analysis and interpretation, writing of the manuscript. LL: Experimental design, data acquisition, analysis and interpretation, writing of the manuscript. DdH: Data acquisition, analysis and interpretation. AA: Data acquisition, analysis and interpretation. TH: Conceptualization, experimental design, data acquisition, analysis and interpretation, writing of the manuscript. JG: Conceptualization, experimental design, data acquisition, analysis and interpretation, writing of the manuscript. All authors edited and approved the manuscript

## Declaration of funding

This work was supported by funding from the Leenaards Foundation (grant 5154.5) to J.G. and a FESSH Basic Science grant to M.M. The study sponsors had no involvement in study design; collection, analysis, and interpretation of data; writing of the manuscript; or the decision to submit the manuscript for publication.

## Conflict of interest statement

No conflict of interest.

## Ethical approval information

The study protocol was approved by the Ethic Committee of Canton Vaud (CER-VD 2024-0117).

## Data availability

All data supporting the findings of this study are available within the Article and its Supplementary files, or are available from the corresponding author upon reasonable request.

## Notes

### Competing Interest Statement

The authors have declared no competing interest.

