## Supplementary Material for "Subchondral bone marrow adipose tissue lipolysis regulates bone formation in hand osteoarthritis"

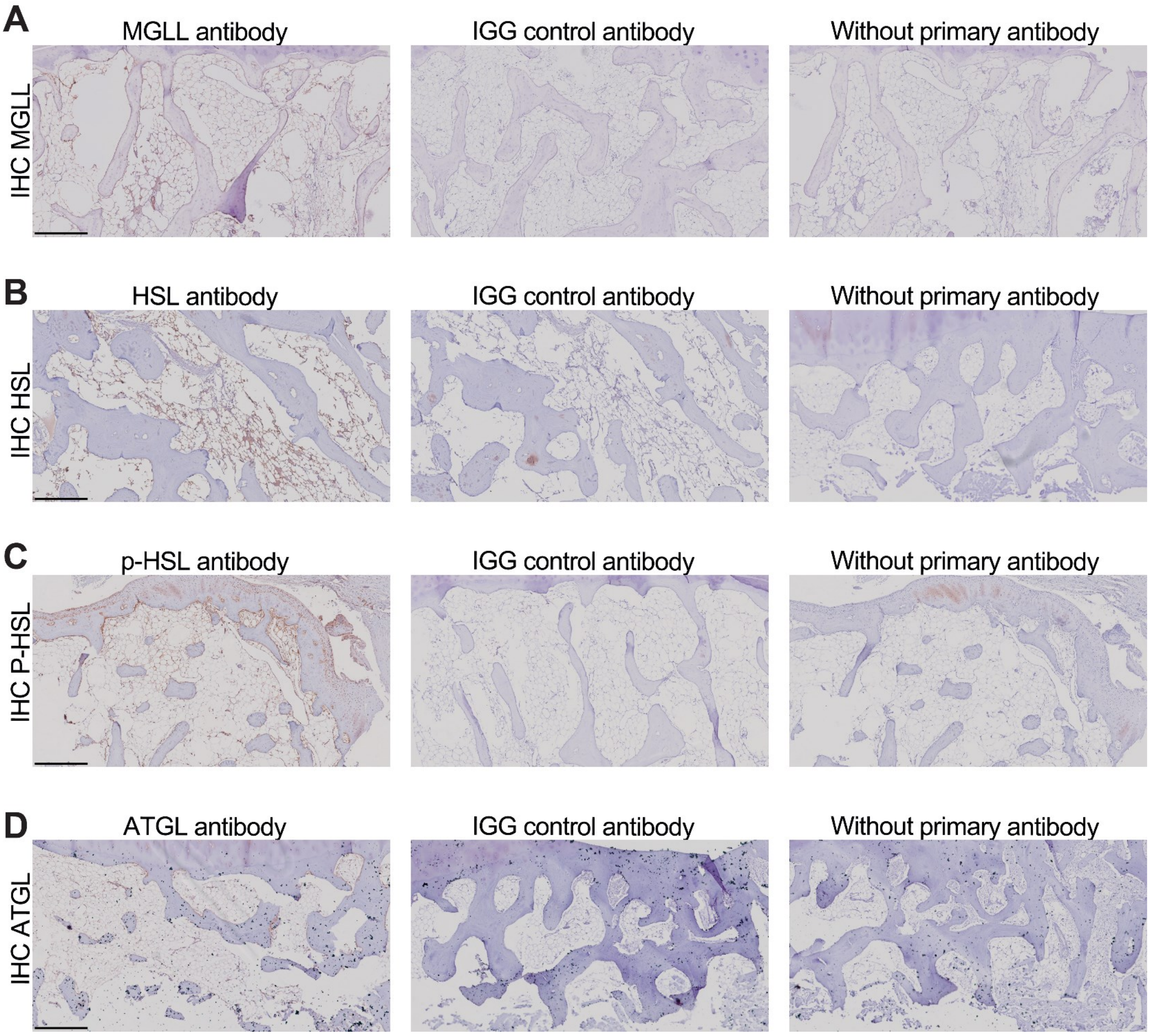


**Figure S1.** Negative controls for immunostaining of MGLL (A), HSL (B), phospho-HSL (C) and ATGL (D) including isotype (middle) and secondary antibody only controls (right).

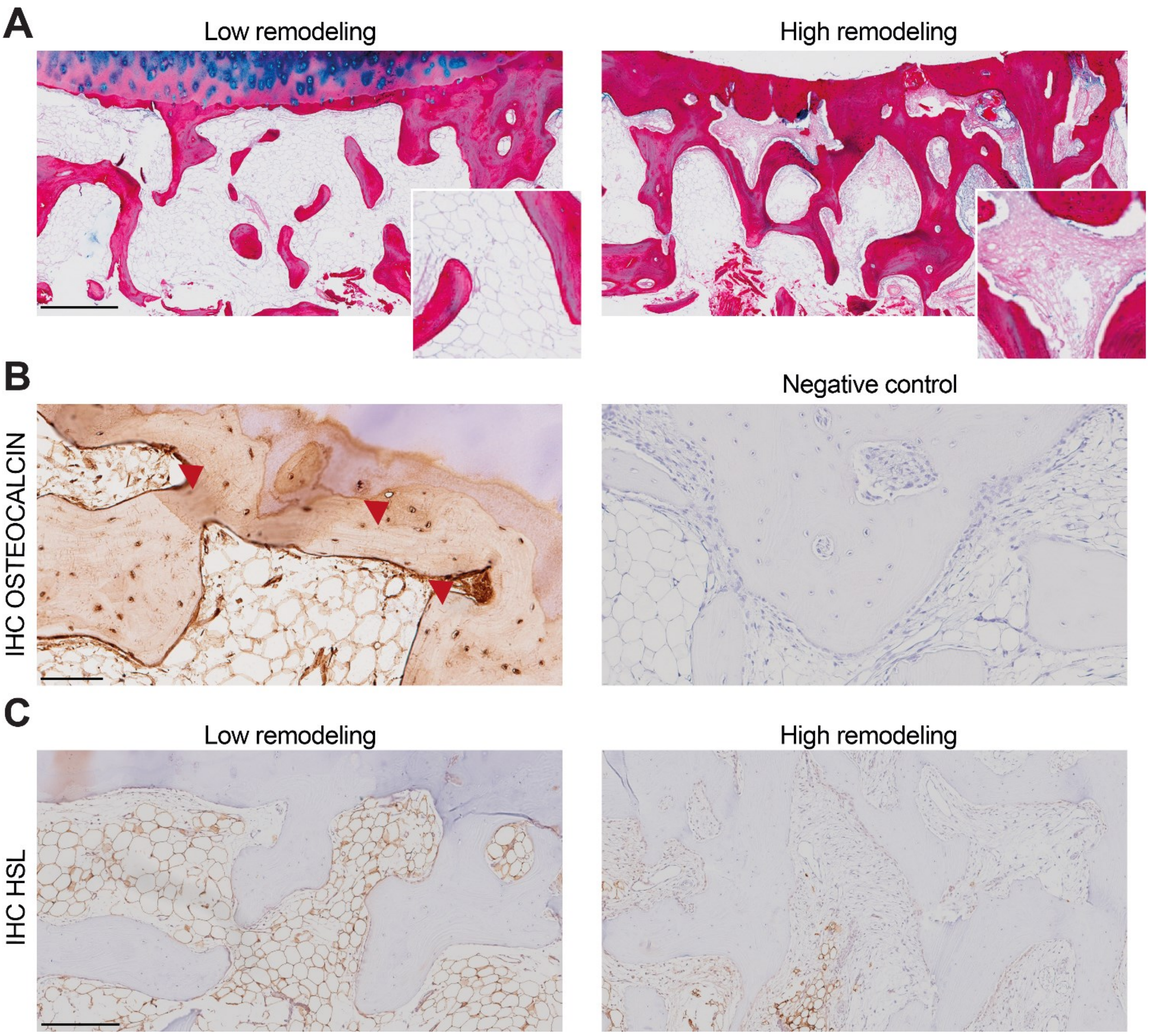


**Figure S2.** Subchondral BMAT remodeling including bone formation and fibrosis visualized by fibrosis visualized by Picrosirius/Alcian blue/Fast Green staining. (A) Osteocalcin immunostaining with osteocytes and osteoblast marked with arrowheads. (B) Representative image of HSL immunostaining marking BMAds in low and high remodeling areas. (C)

**Table S1.** Primer sequences

| *Gene symbol* Accession no. | Forward primer | Reverse primer | Amplicon (length/efficiency) |
|---|---|---|---|
| *GAPDH* NM_002046.7 | tcctgcaccaccaactgcttag | tggactgtggtcatgagtccttc | 86bp / 104.2% |
| *MGLL* NM_007283.7 | ggcatggtactcatttcgcctc | gtttggcagcacaaggttgagc | 99bp /117.9% |
| *PNPLA2* NM_020376.4 | cccacttcaactccaaggac | gcaggttgtctgaaatgccacc | 132bp / 99.3% |
| *LIPE* NM_005357.4 | agccttctggaacatcaccgag | tcggcagtcagtggcatctcaa | 126bp / 123% |
